# Competing mechanisms for the buckling of an epithelial monolayer identified using multicellular simulation

**DOI:** 10.1101/2024.04.01.587527

**Authors:** Phillip J. Brown, J. Edward F. Green, Benjamin J. Binder, James M. Osborne

**Affiliations:** School of Computer and Mathematical Sciences, University of Adelaide, Adelaide, Australia; School of Mathematics and Statistics, University of Melbourne, Melbourne, Australia

**Keywords:** Epithelial Monolayer, Buckling, Modelling, Rigid Body Multi–Cellular Framework

## Abstract

A model using the rigid body multi–cellular framework (RBMCF) is implemented to investigate the mechanisms of buckling of an epithelial monolayer. Specifically, the deformation of a monolayer of epithelial cells which are attached to a basement membrane and the surrounding stromal tissue. The epithelial monolayer, supporting basement membrane and stromal tissue are modelled using two separate vertex dynamics models (one for the epithelial monolayer layer and one for the basement membrane and stromal tissue combined) and interactions between the two are considered using the RBMCF to ensure biologically realistic interactions. Model simulations are used to investigate the effects of cell–stromal attachment and membrane rigidity on buckling behaviour. We demonstrate that there are two competing modes of buckling, stromal deformation and stromal separation.

**Highlights:**

- A rigid body multi–cellular framework allows for the simulation of an epithelial monolayer which is connected to a basement membrane and surrounding tissue stroma.
- Interaction with basement membrane and tissue stroma allows epithelial cells to migrate forming a confluent monolayer.
- Buckling of monolayer can occur through separation from or deformation of the basement membrane.

## 1. Introduction

Epithelial layers are sheets of tightly packed cells that line the surfaces and cavities of organs, forming an interface with the environment [29]. They can be found in numerous parts of the body, including the skin, the bladder, the kidneys, the digestive tract, and reproductive system [29]. A functioning epithelial layer^1^ is made up of three distinct parts [20] (Figure 1):

**Figure 1:**
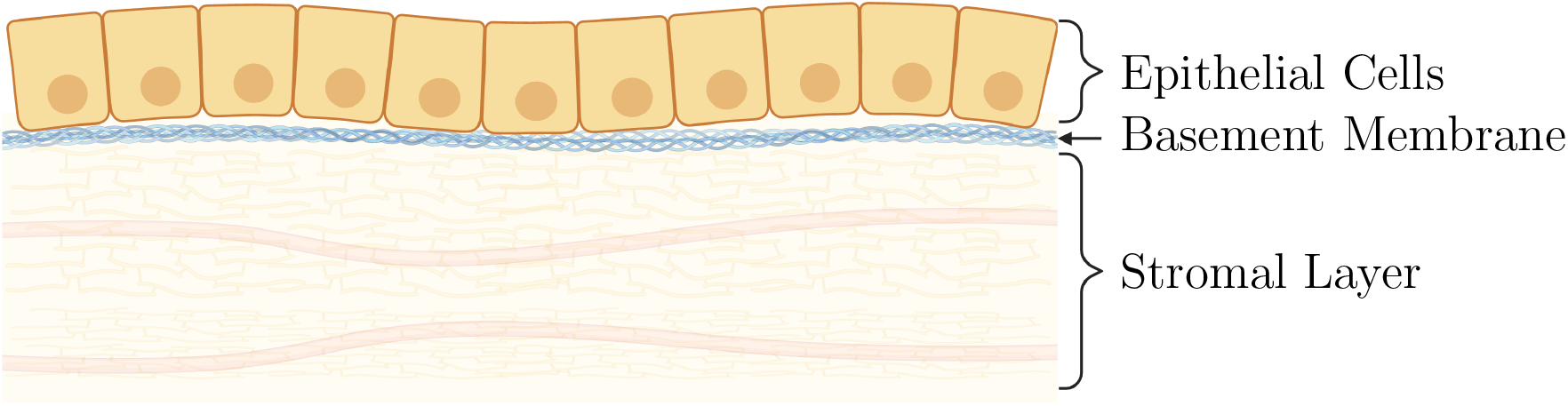
Schematic of epithelial monolayer system. Example of simple epithelial monolayer. Showing the epithelial cells basement membrane and stromal tissue, containing: stromal cells; connective tissue; and blood vessels.

- a stromal layer, made up of connective tissue, vessels and nerves which forms the structure for the cells to sit upon;
- a thin basement membrane layer made up of a web of collagen and glycoprotein molecules which separates the stroma from the cell; and,
- the epithelial cells themselves.

Blood vessels found in the stromal layer transport nutrients to (and in many cases, from) the epithelial cells via diffusion through the connective tissue and basement membrane [29]. The basement membrane provides a substructure that the cells attach to, which persists even when cells are lost. As long as the membrane remains intact, epithelial layers can regenerate [29]. The epithelial cells themselves come in many different types, depending on the particular function the epithelium needs to perform.

There are two broad types of epithelium: stratified and simple [29]. A stratified epithelium is made up of multiple layers of cells, where the bottom-most layer is attached to a basement membrane, and the higher layers sit on top of other cells. Cells proliferate in the bottom layer and transit up through the layers until they die and are shed into the environment [22]. A simple epithelium on the other hand, is characterised by a monolayer of cells where each cell maintains connection to the basement membrane [29]. The single layer functions as a means to absorb and filter material from the environment into the organ [36], as well as secrete material (mucus, enzymes) into the environment [7]. Unlike the stratified epithelium, cells that lose connection to the basement membrane will normally die through a process called anoikis [21]. Loss of this behaviour is understood to be a contributing factor to cancer [21, 24]. By definition, cells in a simple epithelium maintain a single layer, so dividing cells naturally create migratory pressures parallel to the basement membrane [16].

The growth, development, maintenance and regeneration of epithelial tissues are all complex processes involving interactions between cell proliferation and the mechanics of the: cells; membrane; and stroma, and their adhesions to each other. Mathematical and computational models provide a means of integrating a range of experimental observations to bring greater understanding of how this complex web of interactions gives rise to observed behaviours. Both continuum mechanical (*e.g*. elastic) and individual cell based modelling approaches have frequently been used (either by themselves or in combination). However, the relatively simple structure and geometry of many epithelial tissues facilitate their representation using multicellular modelling techniques where individual cells and their interactions are considered. For example, [31] used combination of experiments, a vertex model and a continuum model to understand how cytoskeletal changes in transformed cells interact with the existing tissue geometry to shape the morphology of tumours in the pancreas of mice. The buckling of an ephithelium growing under spherical confinement was investigated by Trushko *et al*. using a similar combination of experiments (using MDCK-II cells), and both vertex-based and continuum modelling [42].

The intestinal epithelium, which is the lining of the small and large intestines, attracts a lot of research attention, in part owing to its association with bowel cancer. This columnal epithelium is comprised of a vast number test–tube–shaped indentations known as the crypts of Lieberkühn [25]. A loss of proper function of the epithelial self renewal process, in these crypts, can lead to pre–cancerous lesions known as polyps, which left untreated can become malignant [6]. Modelling has been used to examine a range of questions about the crypt. One example is clonal conversion which has important applications for carcinogenesis, as it helps explain how a single genetic mutation can spread [43]. Other modelling studies focus on the developmental perspective, for example, aiming to understand how crypt-like structures are formed in colonic organoids grown *in vitro* [4]. The imaginal wing disc of the *Drosophila melanogaster* (fruit-fly) larva is similarly the focus of considerable research interest. In the early stages of growth, a feature appears from the ectoderm made up of cuboidal epithelial cells, which flattens into a disc shape, and through a sequence of changes, eventually becomes a fully formed wing [10]. There are many questions surrounding the processes that regulate the changes, some of which have been investigated using vertex models [19]. Among the questions of interest are: how organ size is regulated during growth [2], how cell growth is regulated by mechanical forces [1], and how the final cell topology is formed [17].

A common modelling approach is to treat the surrounding tissue as static and treat the monolayer as a 2D planar sheet [43] or 2D surface [14, 15]. This kind of modelling is useful to examine things such as clonal conversion, and cell proliferation, however it has limitations due to the fixed structure. An alternative view is to consider a deformable membrane. The common way to achieve this take a cross-section of the layer, representing it as a single row of cells. This approach is best suited to examining movements that happen perpendicular to the layer, including buckling, and cells being ejected from the layer [4, 11, 12]. There are examples where deformable substrates have been implemented in three dimensions [9], however due to their node centric implementation, monolayer deformation is limited [8].

Continuum modelling has also been used to examine the buckling of epithelial layers. The use of such models relies on being able to approximate the tissue as a continuous medium. This can be an unrealistic assumption at certain scales and in certain situations depending on the behaviours that are of interest, but when the conditions are right, it can provide valuable insights. In [16], Edwards and Chapman investigated a flat beam of cells viscoelastically attached to a supporting stromal layer. They see that each of: the applied force; level of cell attatchment; and cell proliferation can independently trigger buckling. Jones and Chapman [26] employ shell theory to explore the role of apical constriction of epithelial cells in gastrulation during embryonic development. Additionally, Nelson and colleagues published two studies that examine the buckling of a one-dimensional beam into crypt-like structures [33], as well as the patterns created by the buckling of a two-dimensional epithelial sheet [34]. The formation of the looped pattern of the gut was considered by Savin *et al*. who showed used scaling arguments based on continuum mechanics that this pattern arises as the result of differential growth between the gut tube and the dorsal mesentery, and their differing mechanical properties [37]. More recent work by Tozluoglu *et al*. used a finite element elastic model to consider folding in the wing disc of Drosophila. Their model, which focused on the roles of non-uniform growth and mechanical properties was able to demonstrate that these factors are key in defining the positions of epithelial folds of the wing disc [41]. In a similar vein, Wyatt *et al*. used a pre-tensioned viscoelastic model to explain the fact that stable folds only emerge in suspended epithelial monolayers of MDCK cells when a ‘buckling threshold’ is exceeded [44].

However, none of these models have been able to model the epithelial layer, and the supporting sub–structure as separate objects, allowing an investigation of their interactions. The Rigid Body Multi–Cellular Framework (RBMCF), from [8], makes this possible. The RBMCF provides us with a range of modelling capabilities, including the ability to represent interacting deformable surfaces. Broadly speaking the RBMCF represents cells, and membranes, as collections of edges, which exert forces on each other. The RBMCF has been used to simulate: a tumour spheroid with decognal cells; a non–intersecting epithelial layer consisting of connected quadrilateral cells; rod based bacterial cells; and spherical cells proliferating in a closed polygonal membrane [8]. One of the key advances of the RBMCF is its ability to represent interactions between surfaces in a mechanically consistent way.

In this paper we exploit the RBMCF to develop a model of an epithelial monolayer, attached to a basement membrane and stromal tissue, in order to understand the mechanisms behind buckling of the epithelial monolayers.

The remainder of this paper is structured as follows. First in Section 2 we present our model of an attatched epithelial monolayer. In Section 3 we present simulations of our model and use it to demonstrate multiple mechanisms for buckling. Finally in Section 4 we discuss the impact of our results and present some avenues for future work.

## 2. Methods

Our model couples the edge based epithelial model (with connected quadrilateral cells) from [8] with an edge based model for the basement membrane and tissue stroma. In the RBMCF, vertices in the model, **x**_*i*_ (for *i* = 1, …, *N*, where *N* is the total number of vertices), move based on the balance of forces due to: the internal model for the cell (or other structures), 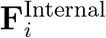; the interactions between cells or between cells and other structures, 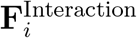; extrinsic noise, 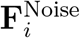; and the viscous drag on the vertices, 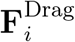. I.e.

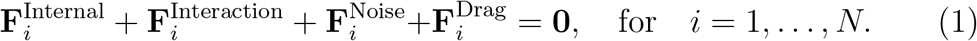

The model equations presented in this study are all non–dimensionalised by a typical cell diameter (taken to be 10*µm* [29]) therefore all positions and lengths are given in terms of Cell Diameters (CD).

This section is organised as follows. We present the model for the epithelial layer in Section 2.1 and the model for the basement membrane and tissue stroma in Section 2.2. The coupling between these models is described in Section 2.3 and, finally, details of model implementation are provided in Section 2.4.

### 2.1. Modelling epithelial monolayer

Following the epithelial layer example in [8] the epithelial layer is defined by a set of connected quadrilaterals (shown in Figure 2 (a)). Let the vertices be denoted by **x**_*i*_ for *i* = 1, …, *N*_*C*_, where *N*_*C*_ is the number of vertices in the epithelial layer. For a given vertex **x**_*i*_, the force due to the embodied energy of the cells is calculated by

**Figure 2:**
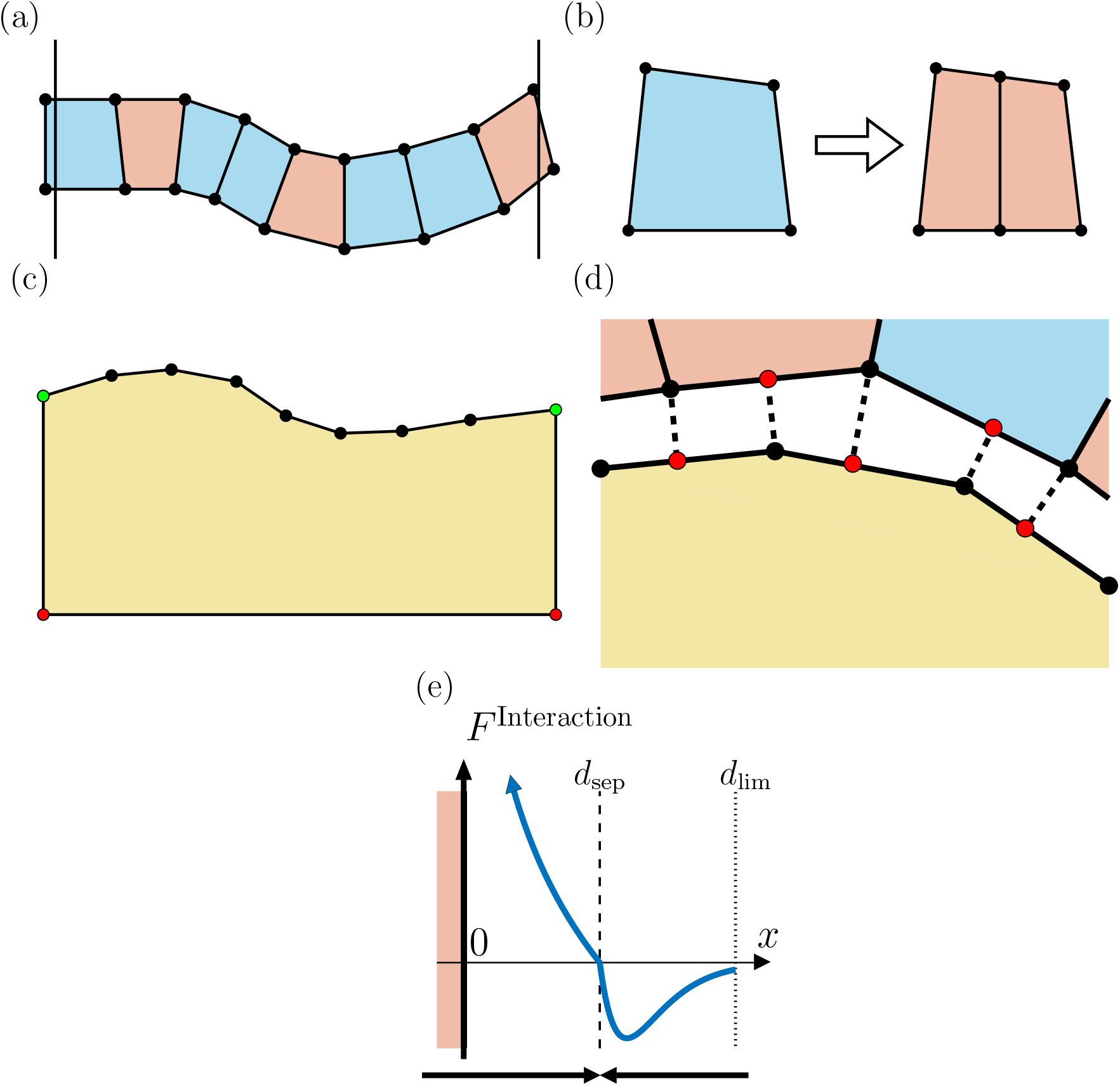
(a) schematic of epithelial monolayer (b) the division process for epitheial cells. When a cell is ready to divide, a new edge is introduced that splits the parent cell into two adjacent daughter cells. The top and bottom edges are split in two by adding a node to their centre. The new edge then joins the new top node to the new bottom node. Modified from [8]. (c) the stromal tissue is represented by a single large object defined by the nodes and edges that make up its boundary. It has a target area and target perimeter and forces are applied to the nodes via energy methods to push it into its preferred shape. The bottom and sides are fixed in place (red and green vertices), while the top surface is free to move (green and black vertices). (d) interaction between the epithelial layer and the stromal tissue. The node from one tissue (in black) interacts with the edges of the other (at the red points). e) schematic of interaction force *F*^Interaction^(*x*). The pink area represents overlapping of cells and stroma (i.e *x <* 0).

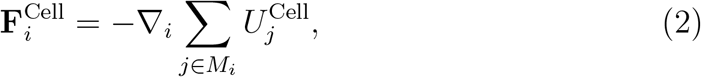

where 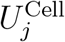 is the energy embodied in cell *j* is, ∇_*i*_ is the gradient with respect to the coordinates of vertex *i* and *M*_*i*_ is the set of all cells that contain vertex *i*.

Following [8] we use energy methods to drive the area and perimeter, of our polygonal cells, in a similar manner to the vertex dynamics model [32]. 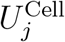 is calculated by

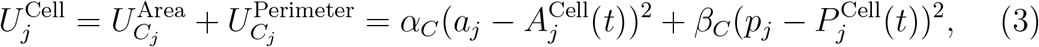

where *a*_*j*_ represents the current area, and 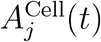 the target area (which changes based on the cell’s age), and where *p*_*j*_, and likewise 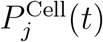, represent the current perimeter and its target value. The parameters *α*_*C*_ and *β*_*C*_ are energy density factors controlling how strongly the cell wants to return to its preferred area and perimeter. Here, following [8], we use the values of *α*_*C*_ = 20 and *β*_*C*_ = 10.

The cell cycle (which controls cell division) is commonly broken up into four phases [3], and occasionally as many as 11 [28]. However, we are mainly concerned with the growth and division processes therefore, following [8], we define two phases: a “growth” phase G where the cell actively increases in size, and a “pause” phase P where the cell maintains its current size. A new cell, immediately after division enters the pause phase where it sits for an amount of time *t*_*P*_ which is drawn from a 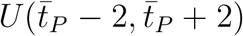 distribution where 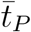 is the mean pause duration (minimum 5^2^). At the end of the P phase, the cell then enters the G phase, where it grows in size over a period of time, *t*_*G*_ drawn from a 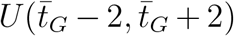 distribution where 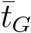 is the mean growth duration (minimum 5). After growth has completed, the cell is able to immediately divide into two child cells by splitting the top and bottom edges in half, and adding a new edge between the new top and bottom nodes see Figure 2 (b).

The actual growth of cell *j* is controlled by setting the target area, 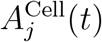, and perimeter, 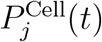, and letting the internal forces cause it to expand. At the end of the growth period the cell will divide into two child cells

Here, we decouple the target perimeter from the target area, and specify their values by the equations

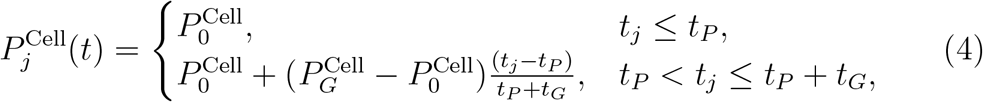

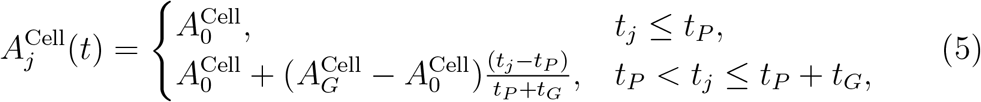

where *t*_*j*_ is the age of cell *j* at time *t*. In this model we set the initial perimeter to *P*_0_ = 3.4, the grown perimeter to *P*_*G*_ = 4, the initial area to *A*_0_ = 0.55, and the grown area *A*_*G*_ = 1. The “grown” values were chosen to have square cells in isolation, and the “initial” values were empirically chosen to minimise pinching^3^, hence preventing artificial layer detachment.

### 2.2. Modelling the basement membrane and tissue stroma

We extend the epithelial ring exemplar from [8], to include the basement membrane and stromal tissues that provides the supporting structure for the epithelial layer. To that end, we model the basement membrane and tissue stroma (referred to here as the stroma) as a large polygonal object, illustrated in Figure 2 (c). The stroma is modelled as a large rectangular body, with the bottom and vertical sides represented by a single edge each, and the top surface made up of multiple edges. The bottom and side edges are fixed in space and the top is allowed to move freely to create a dynamic surface.

Let **x**_*i*_ for *i* = *N*_*C*_ + 1, …, *N*_*C*_ + *N*_*S*_ where *N*_*S*_ is the number of vertices in the stroma.

Internal forces on the stroma are calculated using the same energy methods as used for cells but with one large unconnected polygon. For a given vertex **x**_*i*_, the force due to the embodied energy of the stroma is calculated by

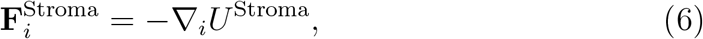

where again ∇_*i*_ is the gradient with respect to the coordinates of vertex *i* and *U*^Stroma^ is the embodied energy of the stroma. A target area and target perimeter are specified, and the difference between those and their measured values is used to calculate the embodied energy:

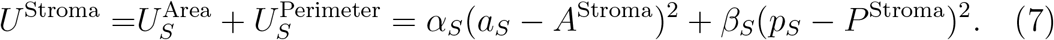

Where *a*_*S*_ (*A*^Stroma^) and *p*_*S*_ (*P*^Stroma^) are the area (target) and perimeter (target) of the stroma respectively. Here the parameters *α*_*S*_ and *β*_*S*_ are the energy factors specifically for the stroma, and they normally will be different from the respective energy factors for the epithelial cells. The target area (*A*^Stroma^ = 39) and perimeter (*B*^Stroma^ = 28.8) will remain constant throughout a simulation, set to the values for the initial rectangular stroma. In this study we will investigate how the stromal mechanical parameters influence the behaviour of the epithelial monolayer.

Note that here, we are assuming that the basement membrane and stromal tissue can be appropriately approximated by an elastic membrane filled with homogeneous fluid. A normal stromal tissue is made up of a wide range of components (blood vessels, ducts, other cells etc.), held together in a connective tissue and future work will use a more complicated model for the stromal tissue.

We fix the bottom vertices of the stroma in place and fix the *x*–coordinate of the upper edge vertices (see the red and green vertices in Figure 2 (c) respectively). Specifically, the edges are held at *x* = 0 and *x* = 10CD. Fixing the edges in place implies a rigid boundary between the tissue of interest and the rest of the biological system. Effectively, this isolates the tissue being modelled from mechanical feedback between it and its surroundings. In various tissues (for example the intestinal crypt), we can see features that may play a similar role to a fixed boundary (e.g. the lamina propria [12]), although in reality, they are not likely to be completely rigid. However, we note that the further away a rigid boundary is placed from where a force is applied, the less impact it should have on the net behaviour. Given the need to maintain the computational tractability of the tissue model, we consider that the approximation of the bottom boundary as rigid is reasonable.

### 2.3. Coupling the epithelium and stroma

To couple the epithelium with the stroma we set the horizontal width of the stroma to 10CD, accommodating an initial population of 20 epithelial cells placed on top of the stroma. We choose this initial cell configuration to represent the compressed nature of the epithelial layer. Epithelial cells divide and push adjacent cells along the stroma and are removed once they extrude past the horizontal boundaries of the stroma, see Figure 2 (a). This is a common approach used for simulating epithelial layers [12].

The epithelial layer will sit on the stroma and interactions between the two will be determined using node-edge interactions (Figure 2 (d)). This allows the layer to remain separate from the stroma, but still affect its shape. Following [8], the interaction force, *F*^Interaction^, between vertices and edges will be calculated according to

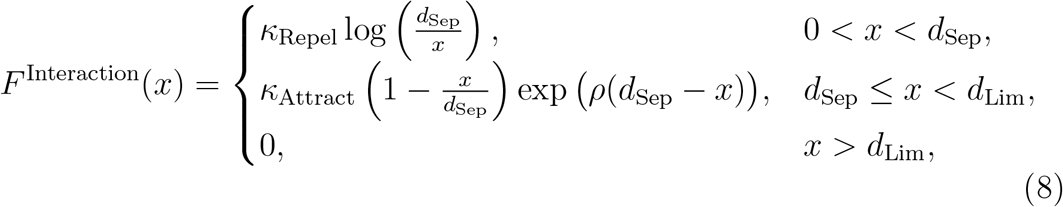

where *x* is the perpendicular distance between edge *k* and vertex *i, κ*_Attract_ is the spring stiffness for attraction (and is zero for cell–cell or stroma–stroma interactions), and *κ*_Repel_ is the spring stiffness for repulsion. Here we choose a fixed value of *κ*_Repel_ = 10 and vary *κ*_Attract_. The typical interaction separation and the maximum interaction distance are given as *d*_Sep_ = 0.1 and *d*_Lim_ = 0.2 respectively. The shape of this interaction force is shown in Figure 2 (e). This form of interaction force has been widely used to model interactions between cells as it allows a variety of interaction distances and magnitudes to be represented [35].

In the analysis to follow, we will only be examining cases up until the point of buckling, meaning that interactions between epithelial cells or between the stroma and itself are unlikely to occur however we include repulsion here in order to avoid overlapping of edges.

When this interaction force acts on the vertices directly (the black circles in Figure 2 (d)) we can calculate the force applied to the vertex directly as

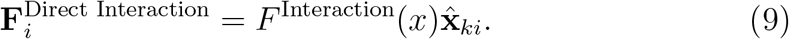

where *x* is the separation between vertex *i* and neighbouring edge *k* (the length of the dashed line in Figure 2 (d)), and 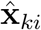 is the unit vector in the direction of the vertex *i* from the point of intersection on the edge *k* (the direction of the dashed line shown in Figure 2 (d)).

When this interaction force acts on an edge *k* (the red circles in Figure 2 (d)), following [8] there are two components, one from the direct interaction with the centre of mass on the edge and one from the perpendicular force which results in a torque on the edge. For a rigid body with a single externally applied force **F** that does not pass through through the centre of drag, it can be shown that **F** can be replaced by a force acting at the centre of drag and a moment [30]. Therefore, following [8], the motion of the body can be completely determined by evaluating the two equations

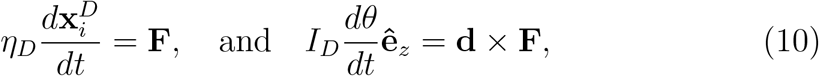

where 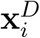 is the centre of drag of the edge, *θ* is the angle the edge makes with the *x*–axis, *η*_*D*_ is the total drag on the edge, *I*_*D*_ is the moment of drag of the edge, and **d** is a vector from the centre of drag to the line of action of **F** such that the two vectors are perpendicular, see [8] for more detail and formal definitions. We can use Equation (10) and a Forward Euler approximation to calculate the motion of the edge due to the applied force **F** in a small timestep *δt*. This can be used to calculate two virtual forces, which when applied to each end of the edge, *k*_1_ and *k*_2_, would give the equivalent motion,

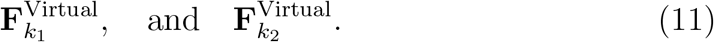

These virtual moves can be combined for each edge that vertex *i* is in to give.

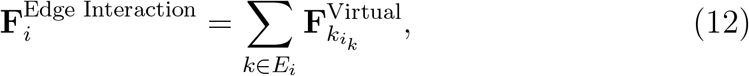

where *E*_*i*_ is the set of edges containing vertex *i* and *i*_*k*_ is the index (1 or 2) that vertex *i* is in edge *k*.

Finally, from Equations (2) and (6), the force applied to the vertex *i* in the model, due to internal interactions, is given by.

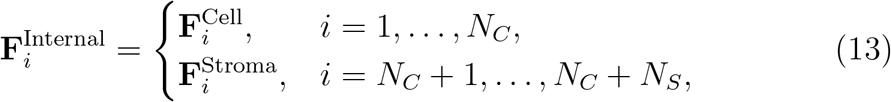

Taking Equations (9), (12) and (13) and balancing the applied force with the drag experienced by the vertex (Equation (1)) allows us to derive the following equations of motion for vertex *i*.

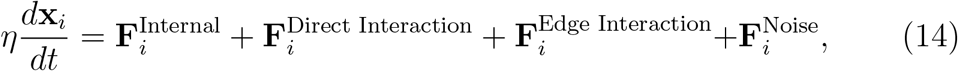

where, following [8], 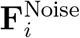 is a randomly directed force which is applied to each node *i*:

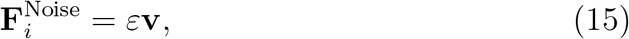

where 0 *< ε* ≪ 1, and **v** is a randomly chosen vector in the unit sphere centered on (0, 0), which is unique at each time step for each node.

We use a Forward Euler approximation for the derivative to simulate the evolution of the vertex locations [8].

### 2.4. Implementation, parameters, and initial conditions

This model has been implemented using the RBMCF modelling tool released as part of Brown et al. [8]. The code is freely available from https://github.com/luckyphill/EdgeBased. Specifically in Release 1.0.2 of the tool.

All parameters used in the model are specified in Table 1. Where possible model parameters are taken from [8]. Those marked with an asterisk are chosen specifically for this study. The stroma specific geometry parameters, *A*^Stroma^ and *P*^Stroma^, are chosen based on the geometry of the epithelial layer simulation. The cell cycle duration parameters, 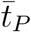 and 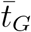, are chosen to cover cell cycle duration’s seen experimentally within epithelial tissues [29].

**Table 1:** Parameter values used in all simulations. The majority of Parameter values are sourced from [8]. The remainder, marked with an asterisk are introduced here and are detailed in Section 2.4. Note, CD=Cell Diameters and CM=Cell Mass.

| Parameter | Description | Value | Units | Source |
| --- | --- | --- | --- | --- |
| $A_0^{\text{Cell}}$ | Cell initial target area | 0.55 | CD<sup>2</sup> | [8] |
| $A_G^{\text{Cell}}$ | Cell grown target area | 1.0 | CD<sup>2</sup> | [8] |
| $P_0^{\text{Cell}}$ | Cell initial target perimeter | 3.4 | CD | [8] |
| $P_G^{\text{Cell}}$ | Cell grown target perimeter | 4.0 | CD | [8] |
| $\alpha_C$ | Cell area deformation coefficient | 20 | CM hours<sup>-2</sup> CD<sup>-2</sup> | [8] |
| $\beta_C$ | Cell membrane tension coefficient | 10 | CM hours<sup>-2</sup> | [8] |
| $A^{\text{Stroma}}$ | Stoma target area | 39 | CD<sup>2</sup> | * |
| $P^{\text{Stroma}}$ | Stroma target perimeter | 28.8 | CD | * |
| $\rho$ | Decay in attractive force | 5.0 | - | [8] |
| $d_{\text{Sep}}$ | Target edge separation | 0.1 | CD | [8] |
| $d_{\text{Lim}}$ | Max interaction distance | 0.2 | CD | [8] |
| $\kappa_{\text{Repel}}$ | Global repulsion magnitude | 10 | CM hours<sup>-2</sup> | [8] |
| $\eta$ | Drag coefficient | 1.0 | CM hours<sup>-1</sup> | [8] |
| $\varepsilon$ | Random noise | 10<sup>-4</sup> | CM hours<sup>-2</sup> | [8] |
| $\bar{t}_P$ | Mean pause duration | 5–12 | hours | * |
| $\bar{t}_G$ | Mean growth duration | 5–12 | hours | * |
| $\alpha_S$ | Stroma area deformation coefficient | 0–20 | CM hours<sup>-2</sup> CD<sup>-2</sup> | * |
| $\beta_S$ | Stroma membrane tension coefficient | 0–20 | CM hours<sup>-2</sup> | * |
| $\kappa_{\text{Attract}}$ | Cell–stroma attraction magnitude | 0–20 | CM hours<sup>-2</sup> | * |

The ranges for the dynamical parameters swept over in this study, *α*_*S*_, *β*_*S*_, and *κ*_Attract_, are chosen to include equivalent parameters used in the model, taken from [8] (e.g., *α*_*S*_ is chosen to be similar to *α*_*C*_).

As discussed in Section 2.3, we start the simulation with 20 connected identical rectangular cells (0.5CD by 1CD) placed *d*_sep_ above a flat rectangular stromal tissue (with initial area *A*^Stroma^ and perimeter *P*^Stroma^). Cells are initialised with a birth time drawn from a 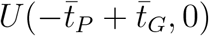 distribution.

## 3. Results

### 3.1. Epithelial layer exhibits buckling

Running the model, we observe different types of behaviour depending on the input parameters. Specifically we set the mechanical parameters *α*_*S*_ = 10, *β*_*S*_ = 6, and *κ*_Attract_ = 10 and vary the proliferation rate. In Figure 3 (a) (and Supplementary Movie 1) we present a simulation where the input parameters intuitively would be conducive to a stable layer (slow proliferation, 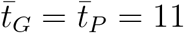), we observe that the epithelium and stroma maintain contact. The layer may exhibit some small deviations, but over long simulation periods, there are no signs of buckling or detachment. In Figure 3 (b) (and Supplementary Movie 2) we present a simulation in the reverse situation (faster proliferation, 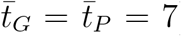). We see buckling and eventually detachment in simulations.

**Figure 3:**
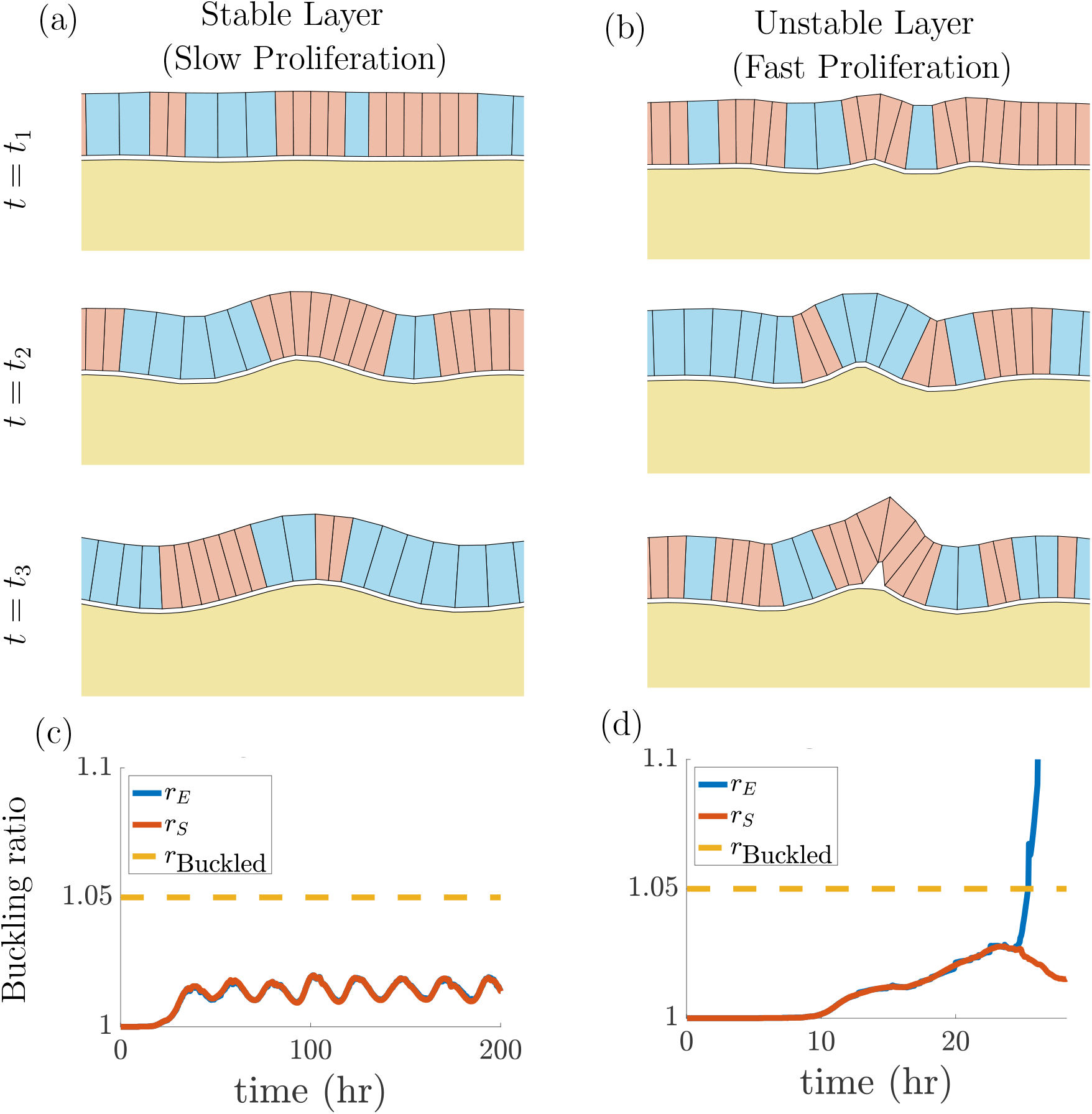
Epithelial monolayer can exhibit buckling. (a) simulation with slow proliferation 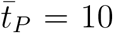. (b) simulation with fast proliferation 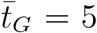. Snapshots are given over time for three indicative times *t*_1_ *< t*_2_ *< t*_3_ (different for (a) and (b)). (c) and (d) show buckling ratios, *r*_*E*_ (blue) and *r*_*S*_ (red), over time for the slow proliferation rate, and (d) for fast proliferation rate. *r*_Buckled_ = 1.05 threshold is shown by the orange dashed line. Mechanical parameters are *α*_*S*_ = 10, *β*_*S*_ = 6, and *κ*_Attract_ = 10. All other parameters as in Table 1. Simulations are stopped at *t* = 200 hours or when *r*_*E*_ = 1.2. Simulations are shown in Supplementary Movies 1 and 2 respectively.

We can examine the causes of buckling systematically by performing a parameter sweep across a range of suitable input values. But first we need a way to determine when buckling has occurred. When an epithelial layer in our model buckles, it goes from having a predictable shape with a bounded number of cells, to some unspecified shape with an exponentially growing hump. A simple way to quantify whether a layer has buckled is to compare the path length of the layer to the value expected for a stable layer. To that end, we define the *buckling ratio*, given by

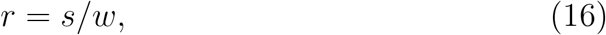

where *s* is the measured path length of the underside of the epithelial cells, and *w* is the horizontal width the layer covers (here 10 CD). By definition, the ratio will be positive and greater than or equal to 1, with values very close to 1 representing a near flat layer, and numbers greater than 1 representing increased undulations in the layer. We can calculate this value for both the epithelial layer *r*_*E*_ and the top surface of the stroma *r*_*S*_.

Buckling ratios are calculated for two sets of simulations (terminated when either *t* = 200 or *r*_*E*_ = 1.2) with slow and fast proliferation rates (see Figure 3(a,c) and (b,d), respectively). In the case of slow proliferation, both buckling ratios are bounded with *r*_*E*_, *r*_*S*_ *<* 1.025 and we observe a stable layer in Figure 3(a,c). However, in the case of fast proliferation only the stroma ratio *r*_*S*_ *<* 1.025 is bounded with the epithelial ratio increasing in time up to the value of *r*_*E*_ = 1.2 used to terminate the simulations (see Figure 3(b,d)). Thus there is an unstable layer in the fast proliferation rate simulations. Based on these observations, we choose *r*_*E*_ = 1.05 to define the point where buckling has occurred as is shown by the dashed orange line in Figures 3 (c,d). Note, the oscillations in Figure 3 (c) are due to oscillations in cell number, which are commonly seen in multicellular simulations (due to the approximate synchronisation of proliferation events) [38]. These increases in cell number create local regions of high compression in the layer which cause it to deform, and then relax as cells move away, causing oscillations in buckling ratio.

### 3.2. Increased proliferation drives buckling

A common observation from simulations of epithelial layers is that increased proliferation can drive buckling [16, 5]. Therefore, in our first experiment, we test buckling over a range of cell cycle lengths by varying the pause phase and growth phase durations. All parameters are fixed to the same value in simulations (See Table 1), except for the values of 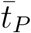 and 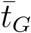 which we vary here. Simulations are run until buckling occurs (defined by *r*_*E*_ ≥ 1.05), or until the simulation reaches 500 hours. Each parameter set is run 50 times, each starting with a different random number generator seed. Note, stochasticity is introduced both through intrinsic noise in cell proliferation and extrinsic noise being added to cell motion.

Figure 4 plots the proportion of buckled simulations for a range of mean pause and growth phase durations between 5 and 12 hours. This range captures interesting features in the parameter space, and is roughly in line with the reported cell cycle lengths for crypt cells in mice [40]. Each point in the plot represents a fixed parameter set where 50 simulations were run. The colour of the point indicates the proportion of those simulations where buckling occurred, with dark red corresponding to all buckling, and dark blue corresponding to all simulations remaining stable for the full 500 hours. Here we can see that faster cell cycles do indeed increase the likelihood of buckling, confirming that the model can reproduce established results. The colours between red and dark blue (i.e. orange, yellow, green, teal) represent parameter sets where some simulations buckled, and some remained unbuckled for the full 500 hours.

**Figure 4:**
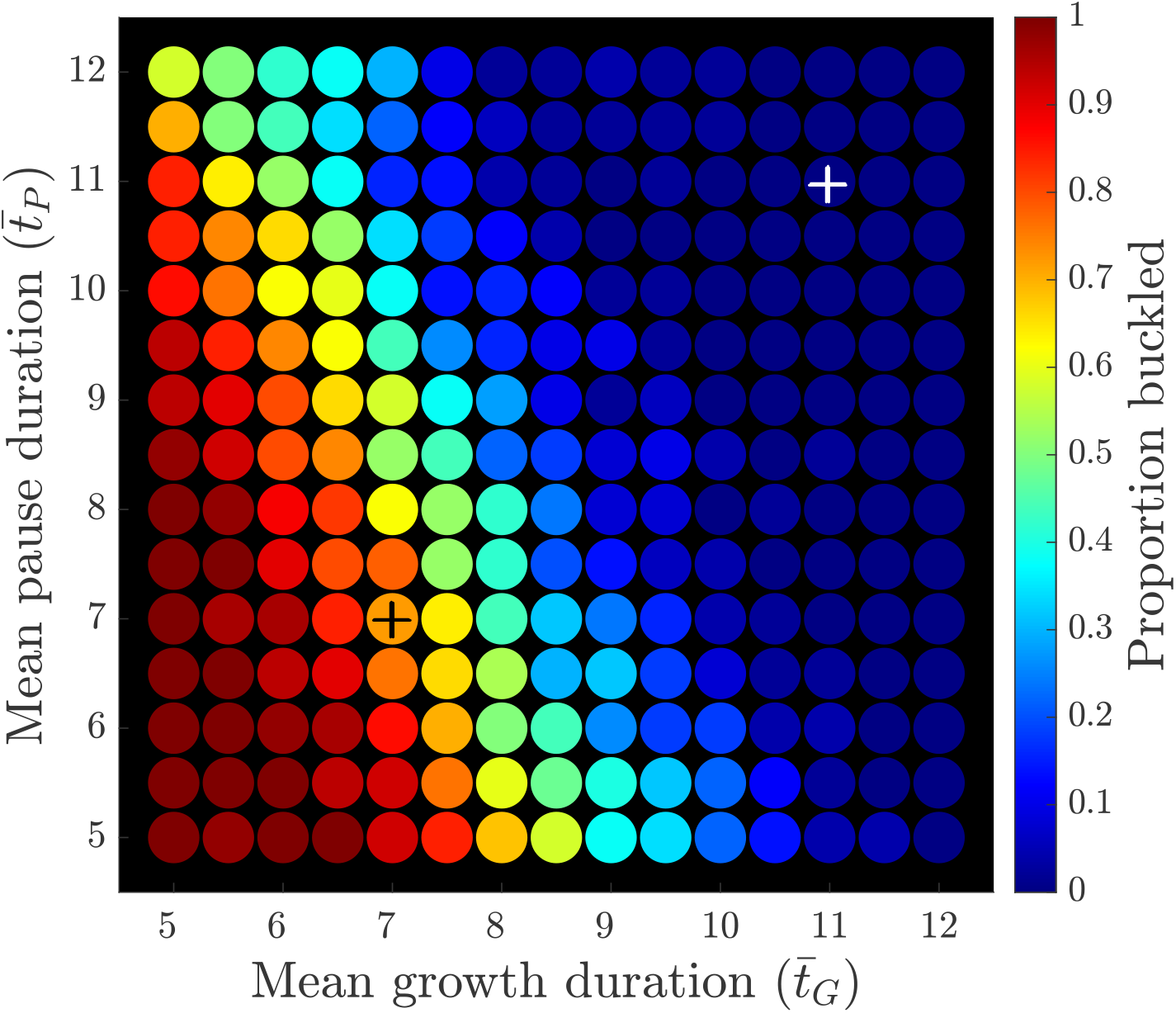
Proliferation drives buckling. Proportion of simulations where buckling occurred for varying values of the pause phase duration and the growth phase duration. Red indicates all 50 simulations exhibited buckling and dark blue indicates all simulations remained stable for 500 hours. As expected, faster cell proliferation (i.e. smaller values of 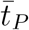 and 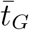) results in a higher chance of buckling. Mechanical parameters are *α*_*S*_ = 10, *β*_*S*_ = 6, and *κ*_Attract_ = 10 All other parameters as in Table 1. Simulations from Figure 3 are shown by the white and black +’s for Figure 3 (a) 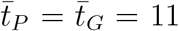 and Figure 3 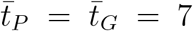 respectively.

In Figure 4 we can observe distinct bands of each colour, forming a “transition front” that can reasonably be fitted with a straight line. If we fit a level curve for parameter sets with specific proportions of buckled simulations, we produce a family of lines with gradient approximately 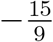. Specifically the level curve for parameter sets where half the simulations buckled forms roughly a straight line with equation 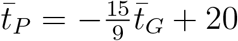. By a simple geometric argument, it is straight forward to see that passing through the transition region from all buckled to none buckled is much faster when 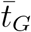 is varied compared to when 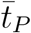 is varied. This implies that changing the length of the growth phase can more rapidly alter the buckling behaviour of the layer. This logically makes sense, since the growth phase is when additional internal stresses are added to the layer, and if these stresses are introduced more rapidly (i.e. with a shorter 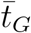), one would expect buckling to be more likely. Reducing the pause phase while keeping the growth phase constant will not change how quickly internal stresses are introduced, but it will alter the total number of cells that are introducing internal stress at the same time. This again, can cause buckling, but as demonstrated, is slower to take effect.

### 3.3. Buckling is independent of stromal compression

A novel feature of this model is its ability to separate the effects of the stromal tissue surface behaviour, the stromal tissue volume behaviour and the strength of adhesion to the surface. We now perform some experiments to investigate how these effects interact with regards to buckling.

As a first experiment, we vary the parameters for area energy, *α*_*S*_, and perimeter energy, *β*_*S*_, for the stromal tissue. The design of the energy method (Equation (7)) comes from the vertex model [18] where the properties of a cell boundary are likened to the surface tension of deforming cell boundaries, and the internal properties are modelled as a compressible elastic body. In that regard, *α*_*S*_ can be thought of as a measure of the tissue’s resistance to compression, while *β*_*S*_ can be thought of as the spring stiffness of the upper surface which is covered by the basement membrane. By sweeping across the two energy parameters, we can investigate the effect of the stromal tissue compared against the effect of the basement membrane (with all other parameters held fixed, as in Table 1, with 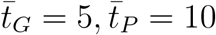, and *κ*_Attract_ = 10).

As with the previous experiment, we run 50 simulations for each distinct parameter set and count the proportion of simulations where buckling occurs (*r*_*E*_ ≥ 1.05) before 500 simulation hours have elapsed.

Figure 5 illustrates the parameter sweep for 1 ≤ *α*_*S*_, *β*_*S*_ ≤ 20, with all other parameters held constant. Here we can see a clear vertical band indicating almost zero effect from varying *α*_*S*_, and a pronounced effect from varying *β*_*S*_ over the same range. This leads us to the conclusion that the compression resistance of the stromal tissue has no impact on the buckling of the epithelial layer.

**Figure 5:**
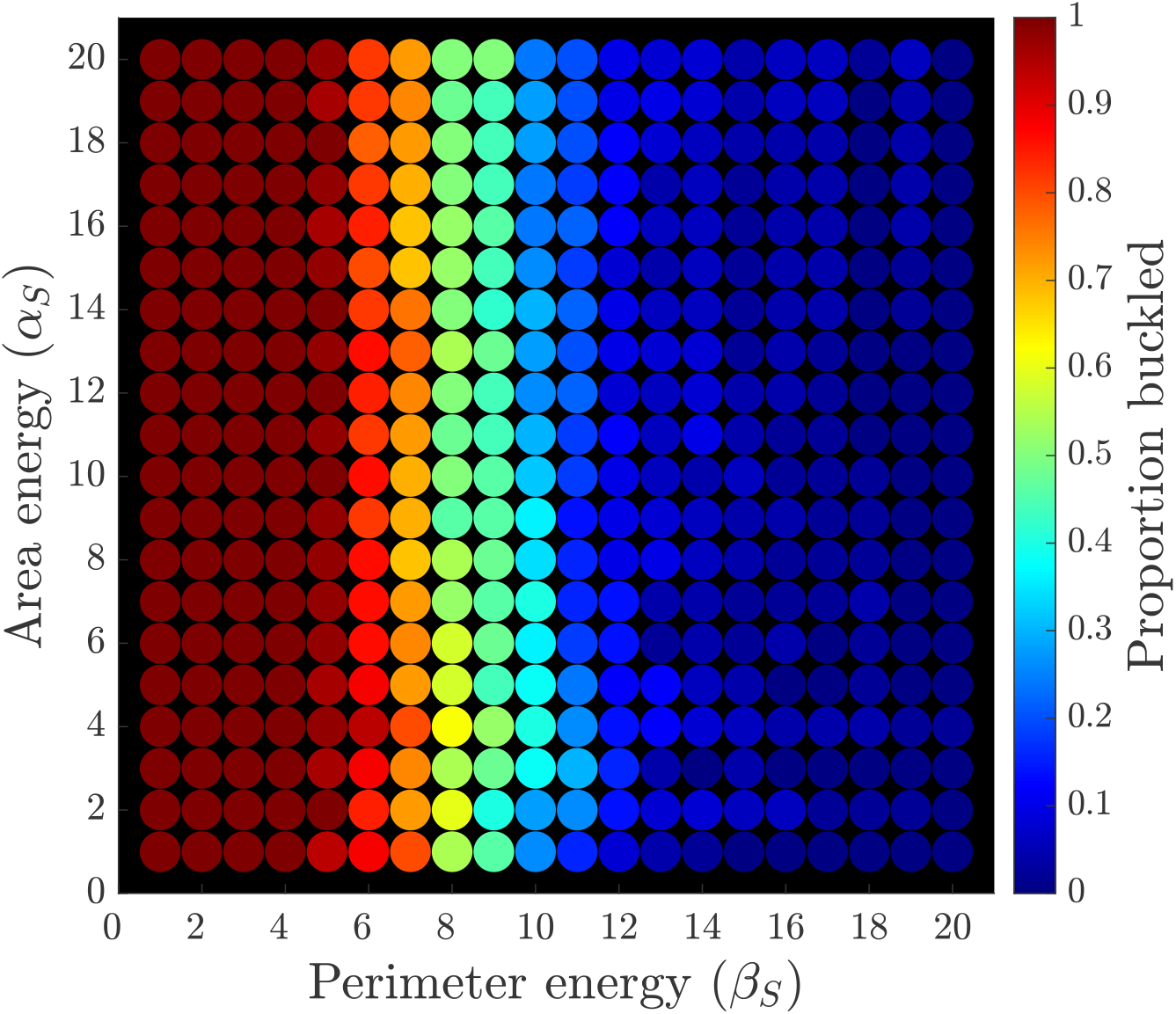
Effect of stromal parameters on buckling. Proportion of simulations where buckling was observed when varying the area energy *α*_*S*_ and the perimeter energy *β*_*S*_. Each point represents 50 simulations, and the colour represents the proportion of those that buckled before the end of 500 simulation hours. 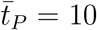, 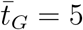, *κ*_Attract_ = 10, and all other parameters as in Table 1.

In the context of the model this is a sensible result. The bottom and side boundaries of the stroma are fixed in place, so the only way the area or perimeter can change is by the top free surface deforming. In its equilibrium state, the surface forms a straight line from corner to corner. Any deviation from this state will lengthen the perimeter, since the shortest distance between two points in Euclidean space is a straight line. However, for a given small deviation from a straight line, it is possible to introduce an additional deviation that restores the area. In other words, there is only one spatial configuration that results in zero perimeter energy, but there are many that result in zero area energy. In a biological context, this model behaviour is unrealistic; stromal tissue is made up of numerous components that exhibit visco-elastic properties as discussed in Section 1. Approximating it to a homogeneous fluid ignores these properties. That said, much of a tissue is made up of fluid, so there will be some degree of comparable behaviour between the model and a physical tissue. To the extent that a real tissue behaves like a fluid, this result suggests the basement membrane influences the chance of buckling for a epithelial monolayer to a much greater degree than the stroma does.

### 3.4. Buckling can occur through multiple mechanisms

We now turn our attention to the interaction force, between the stroma and epithelial layer, which is given by Equation (8). The parameter *κ*_Attract_ captures the strength of attraction between the water-bed stroma and the rectangular cell model of the epithelial layer, hence by increasing or decreasing *κ*_Attract_ we can simulate a stronger or weaker attachment between basement membrane and epithelial cell. Intuitively, a weaker interaction should lead to increased surface detachment and buckling. However, in Figure 5 we also demonstrated that buckling is driven by the surface tension in the membrane, *β*_*S*_. We now perform an experiment to test the interaction *β*_*S*_ and *κ*_Attract_. Once again, we perform a parameter sweep across the two variables keeping all others constant (as in Table 1, with 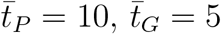, and *α*_*S*_ = 10). We run 50 simulations for each parameter set (each starting from different random seeds), and track the proportion of simulations that end in buckling (*r*_*E*_ ≥ 1.05) before completing 500 simulation hours.

Figure 6 (a) illustrates the parameter sweep for varying stromal stiffness and membrane adhesion (1 ≤ *β*_*S*_, *κ*_Attract_ ≤ 20). As anticipated, the plot demonstrates that both weakening the layer interaction with the surface and lowering the perimeter energy increase the likelihood of buckling. What is not clear however, is whether the buckling in these two cases is caused by the same mechanism. Figures 6 (b)–(c) illustrate snapshots from two different modes observed from simulations. Figures 6 (d)–(e) illustrate the corresponding buckling ratio over time. For high perimeter energy and low adhesion (*β*_*S*_ = 15, *κ*_Attract_ = 5), the stroma remains relatively flat, but when the conditions are right, the epithelial layer buckles and breaks away, Figures 6 (b) and (d). In the opposite case low perimeter energy and high adhesion (*β*_*S*_ = 1.*κ*_Attract_ = 20), buckling occurs while the layer maintains attachment to the stroma, Figures 6 (c) and (e). This buckling is driven by the increased adhesion leading to a higher number of cells exerting a larger force on the stromal layer leading to deformation. This creates regions of high local curvature, which eventually lead to the layer once again pulling away. Buckling can occur in two different ways, either by the layer breaking away from a flat stroma, or losing connectivity at the peak of a local undulation.

**Figure 6:**
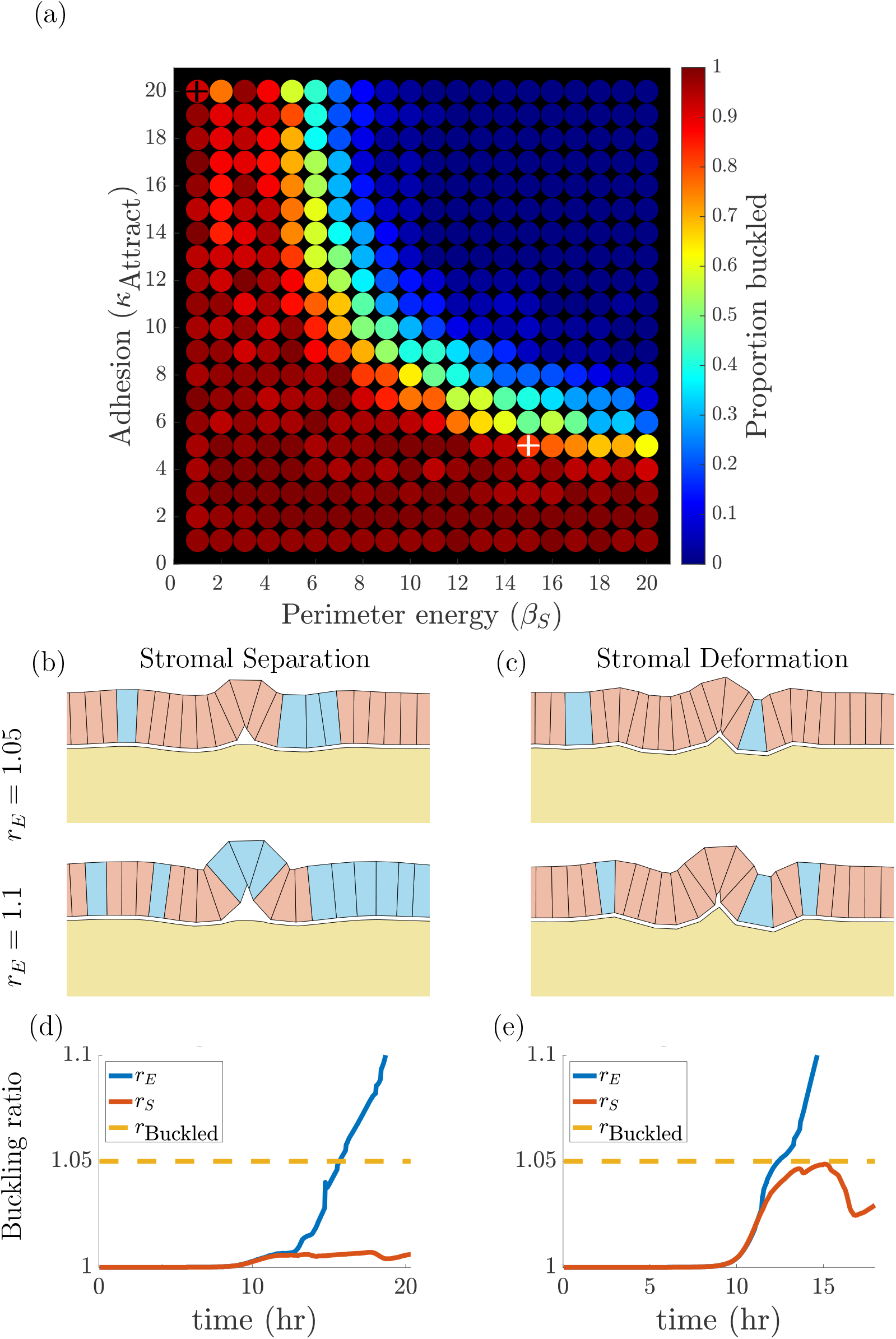
Buckling driven by low membrane adhesion or stromal deformation. (a) plot showing the effect of varying both perimeter energy and and membrane adhesion parameters. Here we can see that buckling can occur for different reasons: if the surface has low stiffness, or if the layer is weakly attached to the membrane. (b) example simulation where high internal pressure in the layer is relieved by the epithelial layer lifting away from the stroma. This situation is driven by a weaker membrane adhesion (low *κ*_Attract_ = 5) with a relatively stiff stroma (high *β*_*S*_ = 15), shown by the white + in (a). The simulation is shown in Supplementary Movie 3. (c) example simulation where weak stroma stiffness (low *β*_*S*_ = 1) leads to the stroma itself buckling into a hump, then a proliferation event near the hump causes the layer to disconnect despite strong membrane adhesion (high *κ*_Attract_ = 20), shown by the black + in (a). The simulation is shown in Supplementary Movie 4. (d) and (e) show the corresponding buckling ratios, *r*_*E*_ (blue) and *r*_*S*_ (red), over time. *r*_Buckled_ = 1.05 threshold is shown by the orange dashed line. Note these simulations are stopped when *r*_*E*_ = 1.2. 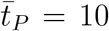, 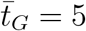, *α*_*S*_ = 10, and all other parameters as in Table 1.

We can investigate where the different modes occur by examining the buckling ratio for the stromal surface *r*_*S*_. We have defined buckling to occur when *r*_*E*_ ≥ 1.05, so we can use the value of *r*_*S*_ at this time to determine which buckling mode has occurred. Looking at Figures 6 (d) and (e) we can see that if *r*_*S*_ ∼ *r*_*E*_ = 1.05, then we know the layer has stayed attached to the stroma up to the point of buckling (Figure 6 (e)), but if *r*_*S*_ *<< r*_*E*_ = 1.05, then detachment has occurred to trigger buckling (Figure 6 (d)).

Figure 7 illustrates the average value of *r*_*S*_ when buckling occurred for each of the parameter sets. Not all repetitions (i.e. random seeds) exhibit buckling for a given parameter set, particularly in the transition region, so the average *r*_*S*_ is calculated only from those simulations that did end by buckling. Red indicates the average stroma buckling ratio is equal to the layer buckling ratio (i.e. *r*_*E*_ ∼ *r*_*S*_), and blue indicates the stroma is close to flat (*r*_*S*_ ∼ 1). We can clearly see that there are two distinct regions in parameter space, roughly divided by the line *κ*_Attract_ = *β*_*S*_. To the bottom right of the plot we see that the stroma has a buckling ratio close to 1, indicating that it has remained flat while the layer has buckled, implying separation. Above the line, we see that the stroma is more buckled as *β*_*S*_ (the stromal perimeter energy) is decreased, or *κ*_Attract_ (the layer adhesion) is increased. These observations fall in line with intuition: a surface that in some sense is less stiff will more easily deform under load, and a stronger adhesion force is more likely to pull a stiff surface with it as it buckles.

**Figure 7:**
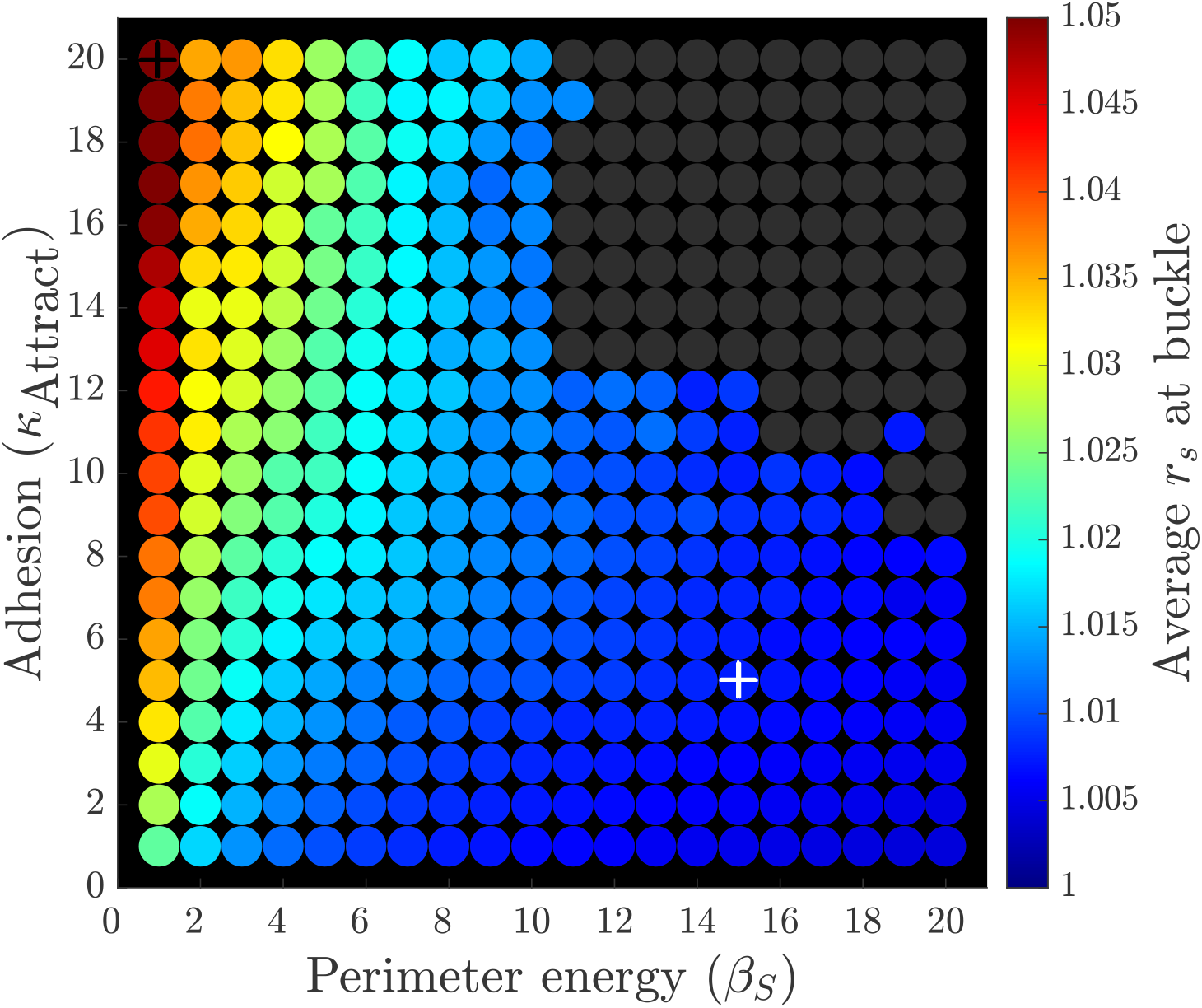
Separation vs deformation. Plot showing the average stromal buckling ratio *r*_*S*_ at the point when we consider the epithelial layer has buckled (*r*_*E*_ = 1.05) for varying surface adhesion and membrane stiffness. For lower adhesion, we can see the stromal surface is close to flat (*r*_*S*_ ∼ 1.00), even though the layer has buckled. Conversely, for lower perimeter energy, *r*_*S*_ approaches the value where the epithelial layer is considered buckled, indicating the layer and stromal surface are the same shape. The dark data points are where no buckling in any of the 50 runs. The white and black +’s represent the simulations from Figures 6 (b) and (c) respectively 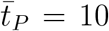, 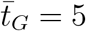, *α*_*S*_ = 10, and all other parameters as in Table 1.

## 4. Discussion

In this study we have applied the RBMCF [8] to a problem that, from a multi–cellular modelling perspective, has seen little attention [5]. This is perhaps due to the difficulty in modelling surface interactions with pre–existing tools. Using our model of a stroma–epithelial interaction, we were able to reproduce the established result that increased proliferation or decreased adhesion is associated with increased buckling [16, 13]. More importantly, the novel construction of the model, specifically including an explicit deformable stroma, allowed us to investigate the impact of basement membrane stiffness and epithelial layer adhesion on the likelihood of buckling. In contrast to existing studies, we were able to show that buckling can occur via two modes: one where the stroma itself is weakened, allowing the layer to buckle while still attached (“deformation”), and the other where internal stresses cause the layer to break away from the basement membrane (“separation”). This result would have not been possible without a modelling approach that can handle two distinct interacting bodies.

The deformation mechanism has been seen in experiments of Drosophila wing disk formation, where a decrease in Extra Cellular Material (ECM), changing stromal stiffness, has led to buckling of the epithelial layer [39]. Trushko *et al*., have created an *in vitro* system of cells proliferating inside a confined region and have demonstrated that the cell layer can detatch causing buckling, as seen in the detachment mechanism in our simulations [42]. Moreover, laser oblation in early stage Drosophila development has been used to show that detachment of the epithelial layer from the surrounding cells can cause buckling [23].

As detailed during the description of the model there are a number of implicit assumptions in the construction of our model impacting how closely it will approximate stromal tissue. We assumed that the basement membrane and stromal tissue can be approximated by an elastic membrane filled with homogeneous fluid. We also considered the system to be represented by a cross sectional geometry. Despite these limitations, the model does allow us a means of investigating how a proliferating monolayer of cells would interact with a deformable supporting structure, and it serves as a good first approximation. Future work will look to work on relaxing these assumptions. First, we will implement an explicit model for the membrane and use a multicellular model (like the spheroid exemplar from [8]) for the surrounding stromal tissue. This will enable us to look at realistic crypt geometries as in [12]. In addition, to look at the processes of crypt budding or fission we could include an evolving stromal area, *A*^Stroma^, or perimeter, *P*^Stroma^. We could also include basement membrane stiffness more explicitly by considering the bending energy of the top stromal surface.

Further work will extend the model to three dimensions, so we have a two dimensional epithelial layer. This will enable us to look at the effect of cell sorting of the heterogeneous cells in the layer which can be seen to induce buckling in, for example, crypt development [27].

## Declaration of competing interest

The authors declare that they have no known competing financial interests or personal relationships that could have appeared to influence the work reported in this paper.

## Data availability

All code to reproduce this study is freely available, under an open source licence, from https://github.com/luckyphill/EdgeBased.

## Acknowledgements

Figure 1 was created with biorender.com. JEFG and BJB’s research is supported by the Australian Research Council (ARC) through DP230100406. JMO’s research is supported by the ARC (DP230100380, FT230100352). This work was supported with supercomputing resources provided by the Phoenix HPC service at the University of Adelaide.

## Author Contributions Statement

PJB conceived and developed the framework, developed the software, ran the simulations, and performed the analysis. PJB, JMO wrote the manuscript. JEFG, BJB and JMO supervised the project. All Authors designed the simulations and analysis. All authors read and approved the final manuscript.

## Supplementary Material

**Supplementary Movie 1** Movie for simulation from Figure 3 (a). Note this runs to *t* = 100 hours instead of 200 as in the figure.

**Supplementary Movie 2** Movie for simulation from Figure 3 (b).

**Supplementary Movie 3** Movie for simulation from Figure 6 (a).

**Supplementary Movie 4** Movie for simulation from Figure 6 (b).

## Footnotes

1 Stromal tissue is not always considered to be part of the epithelial layer. However, we have chosen include it here as the stroma is important in the mechanical behaviour of the epithelial tissue.

2 minimum specified to ensure computational stability.

3 Pinching occurs because of the sudden change of internal area and outer perimeter when a cell divides. This results in unbalanced forces being introduced to the new edge between the two daughter cells, causing it to contract sharply. Pinching can artificially cause layer detachment, but fortunately it can be mitigated by carefully controlling the growth of the cell target area and perimeter.

## Notes

### Competing Interest Statement

JMO EJG and BJB receive funding from the Australian Research Council (ARC).

### Summary of Updates

Edits for clarity. Deeper discussion of results. Linking results to experimental work.

## References

[1] T. Aegerter-Wilmsen, A. C. Smith, A. J. Christen, C. M. Aegerter, E. Hafen, and K. Basler. Exploring the effects of mechanical feedback on epithelial topology. Development, 137(3):499–506, 2010.

[2] T. Aegerter-Wilmsen, M. B. Heimlicher, A. C. Smith, P. B. de Reuille, R. S. Smith, C. M. Aegerter, and K. Basler. Integrating force-sensing and signaling pathways in a model for the regulation of wing imaginal disc size. Development, 139(17):3221–3231, 2012.

[3] B. Alberts, A. Johnson, J. Lewis, M. Raff, K. Roberts, and P. Walter. Molecular Biology of the Cell. Garland Science, 5th edition, 2002.

[4] A. A. Almet, B. D. Hughes, K. A. Landman, I. S. Näthke, and J. M. Osborne. A multicellular model of intestinal crypt buckling and fission. Bulletin of Mathematical Biology, 80(2):335–359, 2018.

[5] A. A. Almet, P. K. Maini, D. E. Moulton, and H. M. Byrne. Modeling perspectives on the intestinal crypt, a canonical system for growth, mechanics, and remodeling. Current Opinion in Biomedical Engineering, 15:32–39, 2020.

[6] M. Bettington, N. Walker, A. Clouston, I. Brown, B. Leggett, and V. Whitehall. The serrated pathway to colorectal carcinoma: current concepts and challenges. Histopathology, 62(3):367–386, 2013.

[7] G. M. Birchenough, M. E. Johansson, J. K. Gustafsson, J. H. Bergström, and G. Hansson. New developments in goblet cell mucus secretion and function. Mucosal immunology, 8(4):712–719, 2015.

[8] P. J. Brown, J. E. F. Green, B. J. Binder, and J. M. Osborne. A rigid body framework for multi-cellular modelling. Nature Computational Science, 1(11):754–766, 2021.

[9] P. Buske, J. Przybilla, M. Loeffler, N. Sachs, T. Sato, H. Clevers, and J. Galle. On the biomechanics of stem cell niche formation in the gut– modelling growing organoids. The FEBS journal, 279(18):3475–3487, 2012.

[10] M. D. de la Loza and B. Thompson. Forces shaping the drosophila wing. Mechanisms of Development, 144:23–32, 2017.

[11] D. Drasdo. Buckling instabilities of one-layered growing tissues. Physical Review Letters, 84(18):4244, 2000.

[12] S.-J. Dunn, P. L. Appleton, S. A. Nelson, I. S. Näthke, D. J. Gavaghan, and J. M. Osborne. A two-dimensional model of the colonic crypt accounting for the role of the basement membrane and pericryptal fibroblast sheath. PLoS Computational Biology, 8(5):e1002515, 2012.

[13] S.-J. Dunn, A. G. Fletcher, S. J. Chapman, D. J. Gavaghan, and J. M. Osborne. Modelling the role of the basement membrane beneath a growing epithelial monolayer. Journal of Theoretical Biology, 298:82– 91, 2012.

[14] S.-J. Dunn, I. S. Näthke, and J. M. Osborne. Computational models reveal a passive mechanism for cell migration in the crypt. PLoS One, 8(11):e80516, 2013.

[15] S.-J. Dunn, J. Osborne, P. Appleton, and I. Näthke. Combined changes in wnt signaling response and contact inhibition induce altered proliferation in radiation-treated intestinal crypts. Molecular Biology of the Cell, 27(11):1863–1874, 2016.

[16] C. M. Edwards and S. J. Chapman. Biomechanical modelling of colorectal crypt budding and fission. Bulletin of mathematical biology, 69 (6):1927, 2007.

[17] R. Farhadifar, J.-C. Röper, B. Aigouy, S. Eaton, and F. Jülicher. The influence of cell mechanics, cell-cell interactions, and proliferation on epithelial packing. Current Biology, 17(24):2095–2104, 2007.

[18] A. G. Fletcher, J. M. Osborne, P. K. Maini, and D. J. Gavaghan. Implementing vertex dynamics models of cell populations in biology within a consistent computational framework. Progress in Biophysics and Molecular Biology, 113:299–326, 2013. doi: 10.1016/j.pbiomolbio.2013.09.003.

[19] A. G. Fletcher, M. Osterfield, R. E. Baker, and S. Y. Shvartsman. Vertex models of epithelial morphogenesis. Biophysical journal, 106(11):2291– 2304, 2014.

[20] R. I. Freshney and M. G. Freshney. Culture of epithelial cells, volume 195. Wiley Online Library, 2002.

[21] S. M. Frisch. Anoikis. Methods in enzymology, pages 472–479, 2000.

[22] E. Fuchs and C. Byrne. The epidermis: rising to the surface. Current opinion in genetics & development, 4(5):725–736, 1994.

[23] M. Gracia, S. Theis, A. Proag, G. Gay, C. Benassayag, and M. Suzanne. Mechanical impact of epithelialmesenchymal transition on epithelial morphogenesis in drosophila. Nature Communications, 10(1):2951, 2019. doi: 10.1038/s41467-019-10720-0.

[24] D. Hanahan and R. A. Weinberg. Hallmarks of cancer: the next generation. Cell, 144(5):646–674, 2011.

[25] A. Humphries and N. A. Wright. Colonic crypt organization and tumorigenesis. Nature Reviews Cancer, 8(6):415, 2008.

[26] G. W. Jones and S. J. Chapman. Modelling apical constriction in epithelia using elastic shell theory. Biomechanics and Modeling in Mechanobiology, 9(3):247–261, 2010.

[27] A. Langlands, A. Almet, P. Appleton, I. Newton, J. Osborne, and Näthke. Paneth cell-rich regions separated by a cluster of lgr5+ cells initiate crypt fission in the intestinal stem cell niche. PLoS Biology, 14 (6), 2016.

[28] C. P. Leblond and M. El-Alfy. The eleven stages of the cell cycle, with emphasis on the changes in chromosomes and nucleoli during interphase and mitosis. The Anatomical Record: an Official Publication of the American Association of Anatomists, 252(3):426–443, 1998.

[29] E. N. Marieb and K. Hoehn. Human anatomy & physiology. Pearson education, 2007.

[30] J. Meriam and L. Kraige. Engineering Mechanics: Statics. Wiley, 5 edition, 2003.

[31] H. A. Messal, S. Alt, R. M. Ferreira, C. Gribben, V. M.-Y. Wang, C. G. Cotoi, G. Salbreux, and A. Behrens. Tissue curvature and apicobasal mechanical tension imbalance instruct cancer morphogenesis. Nature, 566(7742):126–130, 2019.

[32] T. Nagai and H. Honda. A dynamic cell model for the formation of epithelial tissues. Philosophical Magazine B, 81(7):699–719, 2001.

[33] M. R. Nelson, D. Howard, O. E. Jensen, J. R. King, F. R. Rose, and S. L. Waters. Growth-induced buckling of an epithelial layer. Biomechanics and Modeling in Mechanobiology, 10(6):883–900, 2011.

[34] M. R. Nelson, J. R. King, and O. E. Jensen. Buckling of a growing tissue and the emergence of two-dimensional patterns. Mathematical Biosciences, 246(2):229–241, 2013.

[35] J. M. Osborne, A. G. Fletcher, J. M. Pitt-Francis, P. K. Maini, and D. J. Gavaghan. Comparing individual-based approaches to modelling the self-organization of multicellular tissues. PLoS Computational Biology, 13(2):e1005387, 2017.

[36] D. Powell. Barrier function of epithelia. American Journal of Physiology-Gastrointestinal and Liver Physiology, 241(4):G275–G288, 1981.

[37] T. Savin, N. A. Kurpios, A. E. Shyer, P. Florescu, H. Liang, L. Mahadevan, and C. J. Tabin. On the growth and form of the gut. Nature, 476 (7358):57–62, 2011.

[38] M. Slaymaker, J. Osborne, A. Simpson, and D. Gavaghan. On an infrastructure to support sharing and aggregating pre-and post-publication systems biology research data. Systems and Synthetic Biology, 6(1): 35–49, 2012.

[39] L. Sui, S. Alt, M. Weigert, N. Dye, S. Eaton, F. Jug, E. W. Myers, F. Jülicher, G. Salbreux, and C. Dahmann. Differential lateral and basal tension drive folding of drosophila wing discs through two distinct mechanisms. Nature Communications, 9(1):4620, 2018. doi: 10.1038/s41467-018-06497-3.

[40] J. Sunter, N. Wright, and D. Appleton. Cell population kinetics in the epithelium of the colon of the male rat. Virchows Archiv B, 26(1): 275–287, 1978.

[41] M. Tozluoğlu, M. Duda, N. J. Kirkland, R. Barrientos, J. J. Burden, J. J. Munoz, and Y. Mao. Planar differential growth rates initiate precise fold positions in complex epithelia. Developmental Cell, 51(3):299–312, 2019.

[42] A. Trushko, I. Di Meglio, A. Merzouki, C. Blanch-Mercader, S. Abuhattum, J. Guck, K. Alessandri, P. Nassoy, K. Kruse, B. Chopard, et al. Buckling of an epithelium growing under spherical confinement. Developmental Cell, 54(5):655–668, 2020.

[43] I. M. Van Leeuwen, G. Mirams, A. Walter, A. Fletcher, P. Murray, J. Osborne, S. Varma, S. Young, J. Cooper, B. Doyle, et al. An integrative computational model for intestinal tissue renewal. Cell Proliferation, 42 (5):617–636, 2009.

[44] T. P. Wyatt, J. Fouchard, A. Lisica, N. Khalilgharibi, B. Baum, P. Recho, A. J. Kabla, and G. T. Charras. Actomyosin controls planarity and folding of epithelia in response to compression. Nature materials, 19(1): 109–117, 2020.

